# Applying continuous functional traits to large brown macroalgae: variation across tidal emersion and wave exposure gradients

**DOI:** 10.1101/577825

**Authors:** Laura Cappelatti, Alizée R. L. Mauffrey, John N. Griffin

## Abstract

A functional trait-based approach seeks a general understanding of organism - environment interactions, but, among primary producers, its empirical basis rests on vascular plants. We hypothesised that with increasing intertidal elevation, traits of large brown macroalgae would reflect a resource-acquisition vs. conservation (stress tolerance) trade-off at species and community levels. Across the elevation gradient at four UK sites of varying wave exposure, we: i) screened species’ relevant morphological traits, using principal component analysis to reduce dimensionality; and ii) up-scaled species’ traits using community weighted trait means (CWMs). The first principal component (PC1) strongly related to specific thallus area and thallus dry matter content, representing an acquisition - conservation trade-off. Although species generally shifted to the conservative end of this axis as elevation increased, *mid*-shore *Ascophyllum nodosum* sat at the extreme conservative end. PC2 associated with holdfast ratio, thickness and length, with *A. nodusum* scoring higher than other mid-shore species. CWMs of PC1 decreased with elevation at two sites indicating a shift from ‘fast’ to ‘slow’ ecosystem functioning, but this relationship was disrupted by *A. nodusum* at the sheltered site, and by the up-shore extent of *Laminaria digitata* at the most exposed site. The anomalous traits of *A. nodusum* reflect its unique competitive strategy (slow, persistent growth) in the relatively stressful mid-shore. Seaweed functional traits show promise in linking species’ identities to their strategies and ecosystem contributions. However, because resource conservation traits can be related to competitive as well as stress tolerance strategies, predicting seaweed trait responses to environmental stress gradients is challenging.

## Introduction

Characterising how organisms’ functional traits vary along environmental gradients can reveal species’ ecological strategies (Cornwell and Ackerly 2009), the extent of intraspecific variability (Fajardo and Piper 2011), and the adaptive significance of traits (Kraft et al. 2007). Doing so also elucidates how the environment affects trait values at the community level, and therefore the consequences of spatial and temporal variation in the environment for ecosystem functioning (Suding et al. 2008). However, modern functional ecology, a science based on multiple continuous traits measured at the level of individuals, has largely been focused on vascular plants (e.g., Bloomfield et al. 2018) and rarely applied to other, evolutionarily distant, groups of primary producers such as macroalgae or seaweeds.

Here, we adopt approaches of functional ecology to study ‘large brown macroalgae’ (LBM), a functional group including kelps and rockweeds, that dominate standing biomass along temperate rocky intertidal and shallow subtidal zones worldwide (Dayton 1985). These foundation species are major contributors to near-shore primary productivity, and support diverse ecosystem services including fisheries, coastal protection and carbon storage (e.g., Smale et al. 2013). Classic studies in the rocky intertidal point towards functional differences between species of LBM and show the effects of environmental gradients – especially tidal emersion period (represented height on the shore) and wave exposure (commonly represented by wave fetch) – on their distribution (Southward 1959; Ballantine 1961). Furthermore, recent work illustrates how these factors affect the morphology and fitness of individual species (Hepburn et al. 2007). However, categorisation of species into coarse functional groups remains the usual approach in macroalgal ecology (Steneck and Dethier 1994; Arenas et al. 2006; Gómez and Huovinen 2011), and assemblages of LBM have not previously been investigated using a range of continuous functional traits. Therefore, fundamental questions, such as the extent to which gradients such as shore height and wave exposure are associated with functional traits at individual, species and community levels, have remained unexplored.

Like vascular plants, LBM are sedentary, multicellular and differentiated photosynthesising organisms (Hurd et al. 2014). Accordingly, we expect the functional ecology of vascular plants to inform macroalgal functional ecology. Plant ecological strategies range from resource conservative and stress tolerant, to resource acquisitive and competitive (Grime 1974). Functional traits capture these strategies and the underlying trade-offs from stressful to favourable environments (Díaz et al. 2004). For LBM in the intertidal, tidal emersion represents a dominant stress gradient where these shifts in functional traits should be visible. A stress tolerant, resource conservative strategy (e.g., low specific thallus [‘leaf’] area) should be beneficial at the upper shore, where exposure to air for long periods leads to desiccation and slows photosynthesis (Chapman 1995). Meanwhile, a resource acquisitive strategy (e.g., high specific thallus area) should prevail on the more light-competitive lower shore (Littler and Littler 1980). Following plant ecology, we expect strategies and associated trade-offs to reflect in coordinated changes across functional traits (Díaz et al. 2004). Across shore locations, however, gradients of wave exposure, known to be associated with changes in community structure (Ballantine 1961; Burrows et al. 2008) and individual traits (Cousens 1982; Bäck 1993; Blanchette 1997), may disrupt associations between species’ and community-level trait values and tidal emersion. Finally, variability within species threatens to undermine application of traits at the species level; although plant ecologists have addressed its contribution in many systems (Siefert et al. 2015), how intraspecific compares to interspecific variation in trait space is still not known for macroalgae.

We evaluate variability in the functional traits of LBM across gradients of shore height and wave exposure. We conducted surveys and screened individuals for functional traits related to the resource acquisition - stress tolerance trade-offs, along the intertidal at four sites of varying wave exposure in south Wales (UK). Specifically, we investigated: 1) how species of LBM that span shore heights differ in their position in functional trait space; 2) how this interspecific variability scales to the community level along shore heights at sites of varying wave exposure; and 3) given expected intraspecific variability, the extent to which species maintain functional differences across a wave exposure gradient. We hypothesised that, across shore heights, species traits will reflect adaptations to the emersion period in an overall more conservative, stress tolerant, strategy. Although we examined the robustness of these relationships across sites of varying wave exposure, our main focus was on shore height. Table 1 outlines our predictions for how traits would change with shore height, and the mechanisms underlying this variation.

**Table 1.**
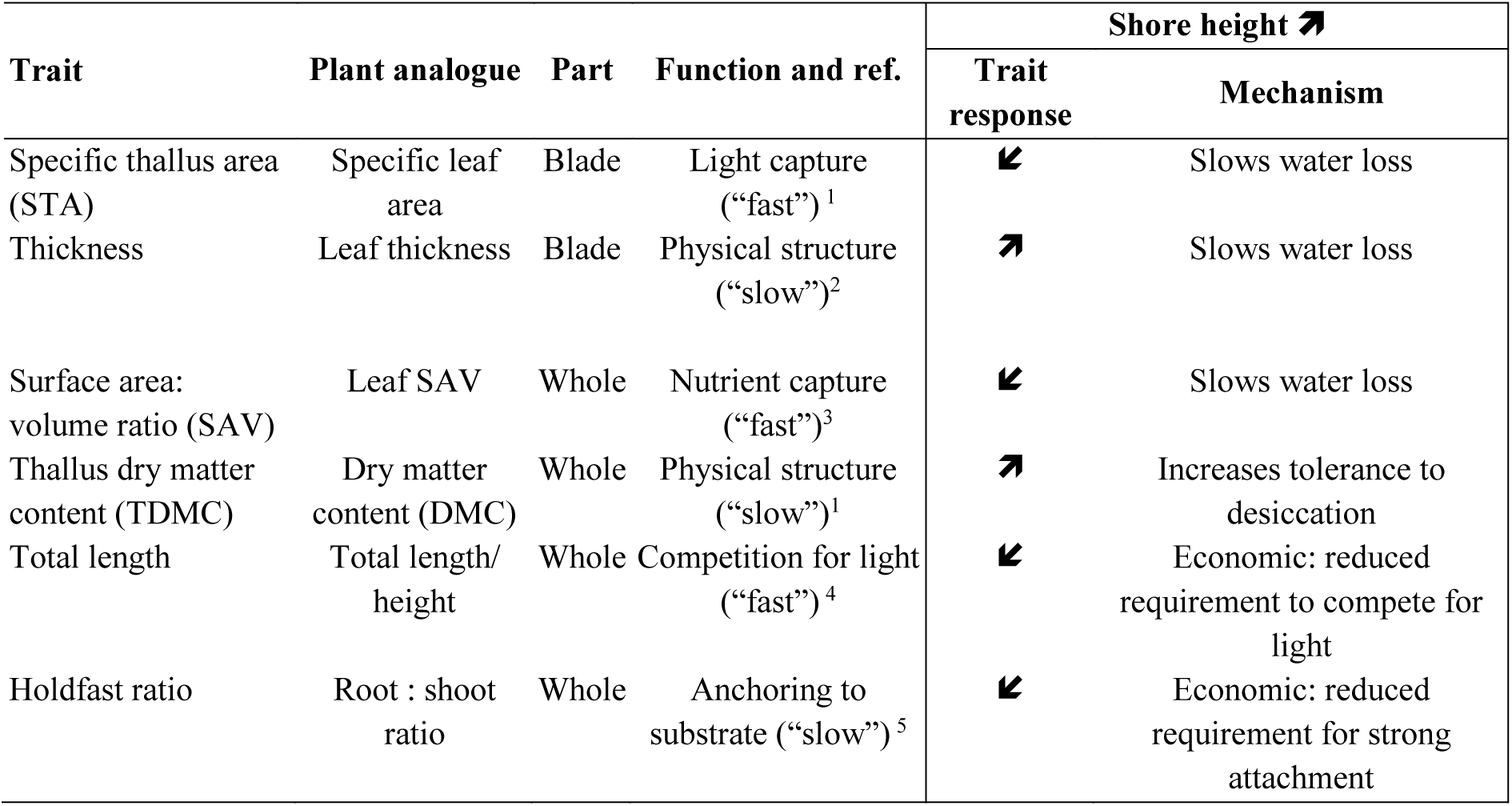
Functional traits measured, their functional relevance, expected variation along the gradients of shore height, and the mechanisms that explain them. Arrows indicate whether trait values are expected to increase, decrease or remain unchanged. “Part” refers to the part of the macroalgae used to measure the trait in this study. “Mechanism” refers to the hypothesised mechanism through which the trait response (rather than the trait itself) is associated with the environmental driver. “Slow” or “fast” indicates the association between the trait and the rate of physiological and ecological processes (see e.g., Reich 2014). Example references supporting each function and shown in superscript numbers are: 1. Wilson et al. (1999); 2. Markager and Sand-Jensen (1996); 3. Littler (1980); 4. Hodgson et al. (1999); 5. Sjøtun and Fredriksen (1995). See *Methods* for explanation of traits and hypothesised responses.

## Methods

### Study sites

The study was conducted on four rocky shores in south Wales (UK), during the summers of 2017 and 2018. Shores in this region experience large tidal ranges, and typically host stands of LBM characteristic of northeastern Atlantic rocky shores. We selected accessible sites covering a wide range of wave exposures and with extensive carboniferous limestone intertidal platforms. Three of the sites span ca. 20 km of the Gower Peninsula (tidal range of 10.37 m): Rhossili (exposed; 51.56 N, 4.32 W), Bracelet Bay (semi-exposed; 51.57 N, 3.98 W), and Oxwich (sheltered; 51.55 N, 4.15 W). Since very sheltered sites could not be found on the Gower, we selected an additional site ca. 50 km west, located in a ria (Milford Haven, Pembrokeshire, tidal range of 7.89 m): Angle Bay (51.68 N, 5.08 W). As an index of wave exposure, we calculated the average wave fetch for each site, which provides good explanatory power for UK rocky shore communities (Burrows 2008; 2012). We used the R packages (R Development Core Team, 2014) *rgdal* (Bivand et al. 2018), to get the spatial points, and *fetchR* (Seers 2017), to obtain the fetch distances of 9 directions, to a maximum distance of 300 km, as default. Average fetch values were 1.95 km at Angle Bay, 16.43 km at Oxwich, 24.79 km at Bracelet Bay, and 67.17 km at Rhossili. We also conducted rapid assessments of water parameters (salinity, temperature, and nutrient levels) and found minimal differences between sites (see Appendix S1 and Table S1 in Supplementary material for details). Accordingly, and in line with a long history of phycological studies (Ballantine 1961; Hepburn et al. 2007), we expected wave exposure to be a key driver of any trait differences between sites.

### Sampling for traits

We assessed morphological traits of LBM at the individual level (within species). Across the intertidal and during low tide, at each site, we haphazardly positioned nine replicate 1 x 1 m quadrats, at least 5 m apart, providing 36 quadrats in total. Individuals were sampled from each replicate quadrat. Due to the effort necessary to measure traits at this scale, we collected a maximum of three samples per species, per quadrat, adding to a total of n = 167 individuals. Samples were stored in bags with seawater, and kept in coolers until returning to the laboratory, where they were immediately transferred to a freezer (−18°C). Sampling took place in summer 2017 (June and July).

### Survey of LBM communities

In order to fully capture the dominant species of intertidal LBM and the assemblages that they form in the study region, we went back to survey the study sites in the summer of 2018 (July-August). A thorough census allowed us to scale traits to the community level and observe how they change with the environmental gradients (tidal height and wave exposure). Along four transects at each shore, spanning the whole intertidal zone, we placed 1 x 1 m quadrats every 10 m. This also allowed us to accurately quantify the mean tidal height distribution of each LBM species for the species-level analyses. Abundance was estimated by counting how many of the 25 string squares inside the quadrat were covered, then transforming that into percentage values (Dethier et al. 1993). Since the study species share similar gross morphologies, cover provides an appropriate, as well as quick, assessment of abundance (Edmunds and Carpenter 2001, Aquilino et al. 2009)

### Tidal height

The position of each quadrat for both surveys was marked using GPS (Garmin 62 ST, Garmin Ltd., Olathe, KS, US) and its height (above chart datum) ascertained using publicly available LIDAR data (lle.gov.wales). To account for differences in tidal range among sites so they are comparable, we used ‘relative height’ (RH) as a measure of the shore height of replicate quadrats, calculated as:

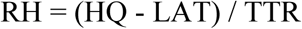

where HQ is the height of the quadrat, LAT is the lowest astronomical tidal height of the specific shore and TTR is the total tidal range of the region (www.ntslf.org). The range of RH spanning the sampled transects varied among sites: Angle bay: 0.22 - 0.70; Oxwich: 0.07 - 0.78; Bracelet Bay: 0.07 to 0.75; and Rhossili: 0.15 to 0.73, possibly owing to differences in exposure, aspect, or shore topography.

### Trait measurements

We assessed six morphological traits that were either known or expected to relate to the ability of individual LBM to cope with environmental stressors of shore height (tidal emersion) and wave exposure (see Table 1). Most of these traits have been measured previously in macroalgae, although not systematically, and have analogue traits that are routinely measured in terrestrial plant ecology (Cornelissen et al. 2003). To allow efficient screening across individuals, the functional traits served as indicators of physiological or physical functions, rather than being direct measurements of them, i.e., ‘functional markers’ (*sensu* Garnier et al. 2004). Before trait measurement, samples were thawed and cleaned from epiphytes, and large fucoid individuals with many fronds were cut by half, longitudinally. Traits were considered at the whole individual or blade-only level (Table 1) and are described below.

The first three traits (1-3) were specific thallus area (STA), surface area to volume ratio (SAV) and thallus thickness. (1) STA and (2) SAV, similarly to specific leaf area (SLA) in plants, capture the ratio between photosynthetically-active surface tissues and structural compounds (Wilson et al. 1999). These traits are expected to be related to (3) thallus thickness, because at constant density a thicker thallus (or leaf) will decrease surface area to mass or volume ratios (Reich et al. 1999). High values of STA or SAV, are associated with higher productivity (Rodriguez et al. 2016; Sakanishi et al. 2017) and nutrient uptake (Hein et al. 1995), but favour water loss and are associated with a weaker physical structure (Littler 1980). Meanwhile, thicker thalli provide structural strength, conferring resistance to frond breakage in wave-exposed conditions (e.g., Wing et al. 2007), but are associated with reduced productivity (Markager and Sand-Jensen 1996) and increase in drag forces. We measured surface area (SA) with software ImageJ (Schneider et al. 2012) from images of samples, cut into thallus parts, spread on an A1 light pad (MiniSun, Manchester, UK). STA was calculated by dividing the blade SA by its dry mass and is expressed as cm^2^ g^−1^. SAV (given as cm^2^ mL^−1^) was obtained from dividing whole SA by total volume, which was measured through water displacement in a graduated cylinder (0.1 mL resolution). For large individuals such as kelp we used a larger cylinder with 1 mL resolution. Thickness is given in cm and was averaged from ten measurements along the blades, using a digital thickness gauge with 0.001 mm precision (Digital Micrometers Ltd., Sheffield, UK).

Three further traits were included: (4) total length, (5) holdfast ratio and (6) thallus dry matter content (TDMC). Total length should determine an individual’s position in the water column, influencing light availability to the thallus (Littler and Littler 1980) and, analogously to plant height (Hodgson et al. 1999), relate to competitive dominance. Total length is the maximum length of the whole individual, and it was measured in centimetres. Holdfast ratio reflects the trade-off macroalgae face between preferentially allocating resources to the fronds to maximise photosynthetic rates and investing in anchoring structures under harsher conditions to prevent dislodgment (Sjøtun and Fredriksen 1995). This trait consists of the ratio of holdfast dry mass to the dry mass of the remaining thallus parts. Dry matter content captures the ratio between structural compounds and water-filled, nutrient-rich photosynthetically-active tissues (Wilson et al. 1999). A high thallus dry matter content (TDMC) should strengthen macroalgae against wave damage and, by reducing the rate of – and tolerance to – water loss, increase desiccation tolerance (Schonbeck and Norton 1979). TDMC was obtained by dividing the dry mass by the fresh mass of the whole individual.

### Statistical analyses

All analyses were performed in R. To explore how LBM species vary in functional trait space, we performed a principal component analysis (PCA) using package *ade4* (Dray and Dufour 2007) on the six traits, using scaled, individual values. Prior to the PCA, we log-transformed the traits and visually checked the linearity of trait-trait relationships. Because of human error during sampling, 58 individual trait measures (representing 6% of all trait values) were absent from our dataset. We therefore conducted analyses with these missing values imputed using the package *mice* (van Buuren and Groothuis-Oudshoorn 2011) and this did not appreciably affect species position relative to each other or trait loadings on axes (see Appendix S2, Table S2, Fig. S2).

To evaluate how this interspecific variability scales to the community level along the shore height gradient and across sites, we examined how community weighted means (CWMs) of loadings from PC1 and PC2 varied with RH at each site. CWM uses species means and assigns proportional weight to the traits according to the abundance of each species, therefore possible to relate to ecosystem functioning, as has been done in plant communities (e.g., Vile et al. 2006). CWMs express the dominant traits in a community (Mokany et al. 2008), and to focus our analyses on communities dominated by LBM we selected quadrats where the total cover of these species was of at least one third (33% of quadrat area). We used the data from the survey and calculated CWMs for each quadrat using package “FD” (Laliberté et al. 2014). Then we plotted the CWMs of each axis for all sites individually to observe their distribution along the relative heights. Because they were not linear, we used Kendall’s rank correlation (which also allows ties) to statistically test the effect of relative heights on traits for each site. We show the data distribution along with pie charts of species’ relative abundances on each point to highlight the species that were driving any trends or patterns.

We next evaluated the degree of intraspecific variability across sites of varying wave exposure, and the extent to which species maintain functional trait differences in face of this variability. We ran a linear mixed model using *lme4* in R (Bates et al. 2015), fitting species by site as predictors of PCs 1 and 2. To account for the possible non-independence of samples collected from the same quadrat, we fitted quadrat identity as a random effect, but this did not improve the model fit (by AIC comparison) and so we report the results of the linear model (ANOVA). Note that we expected the distribution of species to be strongly related to RH due to the obvious zonation, so we did not plan analyses including both species and RH as simultaneous predictors. However, we did test whether residual variance from the above models could be attributed to RH (Appendix S3, Table S3). To integrate multiple traits, we performed the same species by site model using PERMANOVA based on Euclidian distances using *adonis* in the package *vegan* (Oksanen et al. 2018), with the scaled trait scores (Table S4). Two species (*S. latissima* and *H. siliquosa*) were removed from these analyses because they were found at only one site and in less than five quadrats. Of those included, *A. nodusum* and *L. digitata* were both missing from a single site (Oxwich and Angle Bay, respectively); the remaining four species occurred at all four sites.

## Results

We found 8 species belonging to three families. The total number of samples for trait measurement was 14 for *Ascophyllum nodosum*, 27 for *Fucus spiralis*, 45 for *Fucus serratus*, 31 for *Fucus vesiculosus*, 22 for *Pelvetia canaliculata* (family Fucaceae), 3 for *Halidrys siliquosa* (family Sargassaceae), 17 for *Laminaria digitata*, and 8 for *Saccharina latissima* (family Laminariaceae). Species’ mean relative heights and ranges for both surveys are given Table S5.

### The spectrum of functional trait variation

Across sites and shore heights, PC1 captured 46.1% and PC2 captured 21.5% of variation in function trait space (Fig. 1). PC1 most strongly reflected STA, SAV and TDMC and PC2 was associated with holdfast ratio, blade thickness and total length. PC3 captured an additional 16.7% variation and was most associated with length and holdfast ratio (Table S2). Despite intraspecific variability, most species occupied distinct regions of trait space (Fig. 1; Table 2 (ANOVA), species effect: PC 1 and 2: R^2^ = 0.72, P<0.001) and could be associated with traits or axes, illustrating the nature of their functional differences. STA was positively and strongly associated with *S. latissima*, a kelp species found at low shore and restricted to the most sheltered site. Length and holdfast ratio were associated with *L. digitata*, a kelp species of the low shore which is absent from the most sheltered site. TDMC was associated with *P. canaliculata*, the species highest on the shore at all sites. Thickness was associated with the mid shore species *A. nodusum*, which was distinct from other mid shore species *F. vesiculosus* and *F. serratus*. Among the LBM, these *Fucus* species and the upper shore *F. spiralis* shared the most similarity. Trait means and variance for each species are shown on Fig. S1.

**Figure 1.**
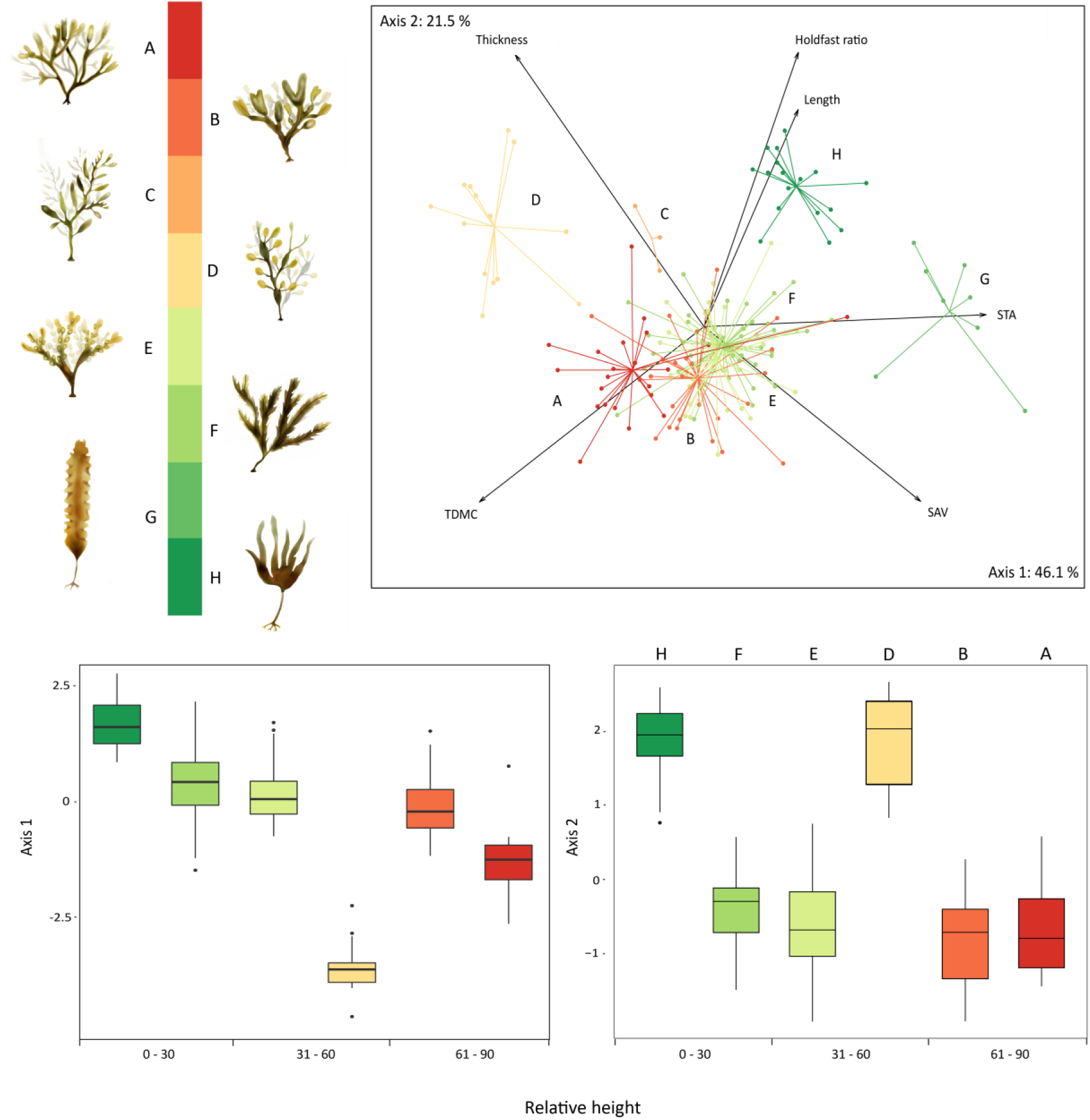
First two PCA axes of large brown macroalgae, based on trait survey from 2017. The top panel shows the distribution of individuals and species in trait space, together with the relationships between functional traits and PC axes. All eight sampled species are included, colour-coded by order of height distribution (top left). Species legend: A, *Pelvetia canaliculata*; B, *Fucus spiralis*; C, *Halidrys siliquosa*; D, *Ascophyllum nodosum*; E, *F. vesiculosus*; F, *F. serratus*; G, *Saccharina latissima*; and H, *Laminaria digitata*. Lower panels show species variation in PC scores, with species sorted by relative height (RH, as proportion, relative to the total tidal variation; full description in *Methods*). Two species were removed from the boxplots due to their low occurrences: *H. siliquosa* and *S. latissima*. Contributions of each trait to the PCA axes are given in Table S2.

**Table 2.** Result of ANOVAs examining the effect of species and site on individual traits loaded onto the first two PCA axes. Degrees of freedom (df), and test statistics (sum of squared differences, mean of squared differences, F-value and P value) are given, as well as an estimated variance explained (Partial R^2^).

|  | df | Sum Sq | Mean Sq | F value | P value | Partial R<sup>2</sup> |
| --- | --- | --- | --- | --- | --- | --- |
| *Axis 1* – R<sup>2</sup>: 0.82 |  |  |  |  |  |  |
| Site | 3 | 30.859 | 10.286 | 20.1006 | <0.001 | 0.080 |
| Species | 5 | 276.483 | 55.297 | 108.0572 | <0.001 | 0.716 |
| Site * Species | 13 | 11.111 | 0.855 | 1.6702 | 0.074 | 0.029 |
| Residuals | 133 | 67.5 | 0.508 |  |  |  |
| *Axis 2* – R<sup>2</sup>: 0.79 |  |  |  |  |  |  |
| Site | 3 | 4.183 | 1.394 | 3.6491 | <0.05 | 0.017 |
| Species | 5 | 176.168 | 35.234 | 92.2094 | <0.001 | 0.719 |
| Site * Species | 13 | 13.372 | 1.029 | 2.7773 | 0.001 | 0.055 |
| Residuals | 134 | 51.202 | 0.382 |  |  |  |

We can observe how species differentiate in PCs 1 and 2 with increasing shore height (Fig. 1, bottom panel). In general, species’ values on both PC1 and PC2 declined with their average shore height. However, *A. nodosum* was a clear outlier, occupying the mid-shore while scoring lower on PC1 than high shore species, and scoring similarly to low shore species on PC2.

### Scaling species’ differences to the community level

The relationships between increasing shore height and CWMs based on PC1 and 2 varied across sites (Figures 2 and 3). PC1 significantly decreased with relative height at the two sites in the middle of the exposure gradient, Bracelet Bay and Oxwich (tau= −0.72 and −0.75, respectively). Accordingly, at these two sites, higher on the shore, dominant species exhibited lower STA and SAV, and higher TDMC. There was also a decrease in PC2 with increasing height at Oxwich (tau= −0.75), indicating that at this site, higher on the shore dominant species were shorter, thicker and with a lower holdfast ratio. There were no clear relationships between height and CWMs of PCs at either the very sheltered Angle Bay or the exposed Rhosilli. At Angle, this can be explained by the dominance and relatively low shore extent of *A. nodusum* (Figures 2 and 3), with its anomalous traits (see above). At Rhosilli, this can be explained by the up-shore extent of the otherwise lower shore *L. digitata*.

**Figure 2.**
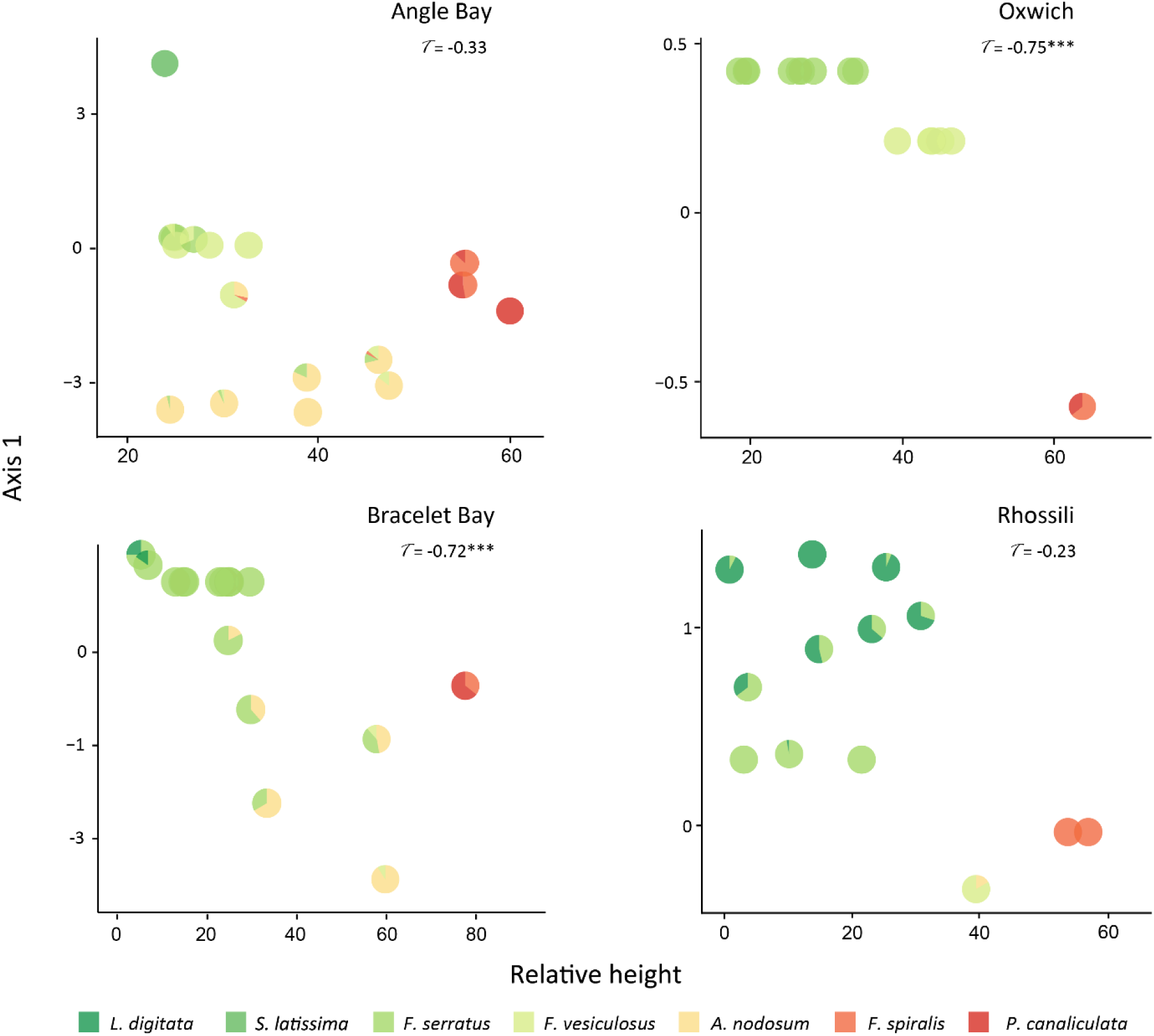
Community weighted means (CWMs) of the first PCA axis along relative heights (as proportion, relative to the total tidal variation; full description in *Methods*. Note that sites differ in range). Pie charts represent species relative abundances for each point (quadrat) and Kendal’s tau is given for each site, with significant values (p<0.01) shown with asterisks.

**Figure 3.**
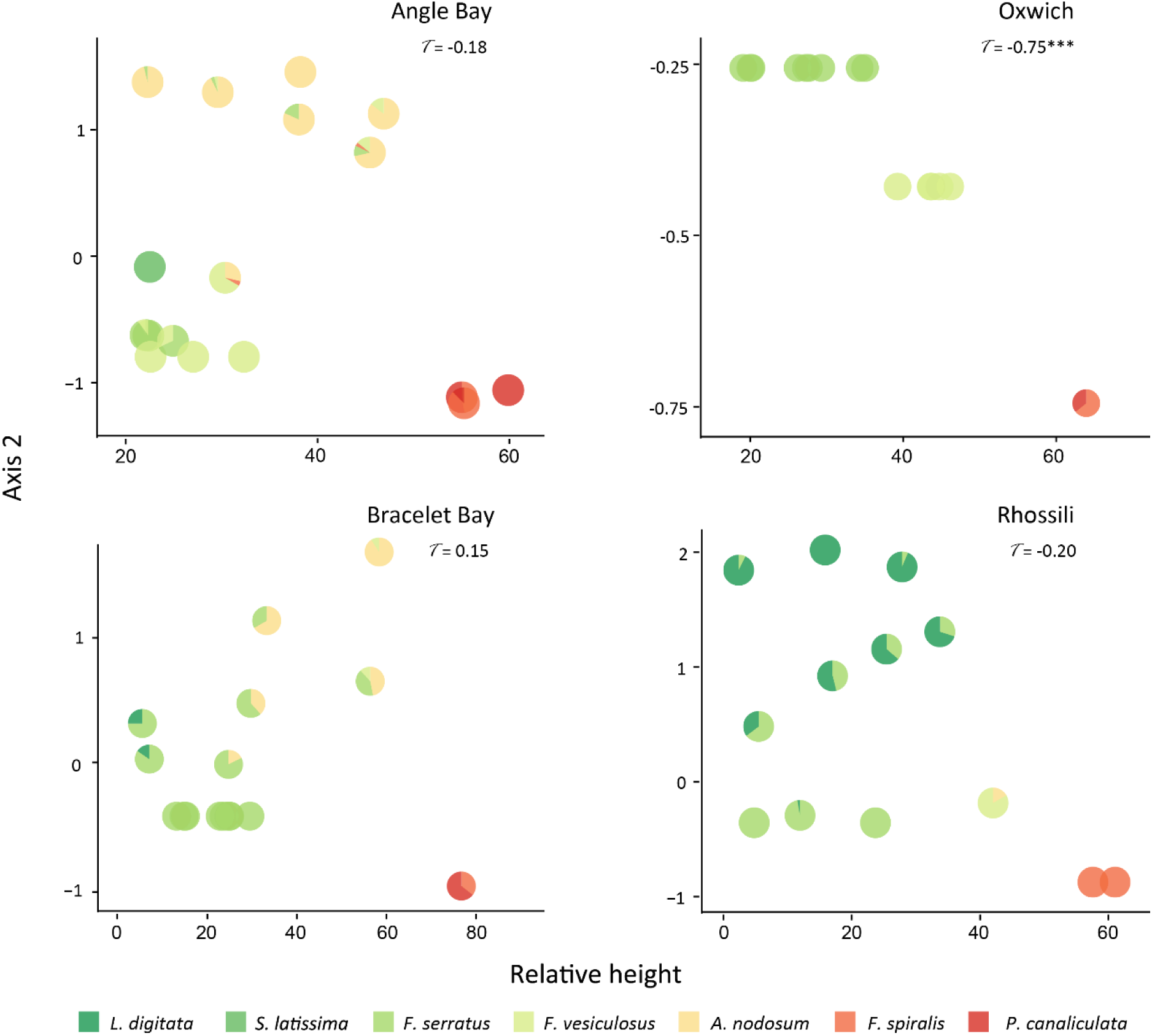
Community weighted means (CWMs) of the second PCA axis along relative heights (as proportion, relative to the total tidal variation; full description in *Methods*. Note that sites differ in range). Pie charts represent species relative abundances for each point (quadrat) and Kendal’s tau is given for each site, with significant values (p<0.01) shown with asterisks.

### Intraspecific and interspecific variability across sites of varying wave exposure

Species’ differences in trait space varied with site. Although there was no clear evidence of a site – species interaction for PC1 (P = 0.07, Table 2) there was an interaction between site and species for PC2 (ANOVA; Table 2). However, these interactions and the main effects of site were much weaker than those of species (PC1: site 8% vs. species 72%; PC2: site 2% vs. species 72%), indicating that the readjustments of species’ relative trait values at different sites are much smaller than the overall differences between them. Residual intraspecific variability could not be attributed to variation in the relative height of collected samples (Appendix S3, Table S3).

## Discussion

Addressing trait variability at both within and between species levels provides insights to inform the study of functional diversity in macroalgal communities. We investigated a group of dominant macroalgae in rocky intertidal shores, the “large brown macroalgae” (LBM) to reveal the levels of morphological trait variability and how these relate to environmental gradients and ecological strategies. We therefore bring a new perspective to the study of macroalgal functional ecology, which has so far remained largely restricted to classifying species into - and comparing - functional groups.

Trait variability among LBM was largely captured by two key axes and could be related to species’ position on the shore and ecological strategies. The existence of two major axes of variation (at least based on the suite of traits measured here) is consistent with studies of vascular plants (Pescador et al. 2015; Díaz et al. 2016). The first, analogous to the ‘leaf economics’ axis in plants, is representative of resource acquisition vs. conservation/stress tolerance trade-offs. That is, LBM will either have high STA and SAV, and therefore high productivity, or have a high TDMC and more resistance to mechanical or desiccation stress. The second axis requires more interpretation, but we argue that it relates to competitive dominance. Longer LBM have higher canopies that are competitively dominant for light, while a greater proportional investment in anchoring holdfasts and thicker blades provide physical resistance to the increased drag associated with larger fronds (Wernberg and Thomsen 2005).

With the exception of *Ascophyllum*, species’ positions on these axes broadly reflect their position on the shore, as we hypothesised, with species higher on the shore scoring lower on PC1 and 2. The outlying traits of the mid-shore *Ascophyllum* appear to reflect its unique strategy among mid-shore species: slow and persistent growth, resisting physical and biological stressors over a long lifespan (10-15 years; Sundene 1973), to eventually dominate late-successional assemblages at sheltered sites (Topinka et al. 1981, Vadas et al. 1990). This strategy appears to require traits that are more conservative than other mid-shore species and also shifts it higher on the second axis reflecting greater thallus thickness (especially) and length. The case of *Ascophyllum* suggests that the first trait dimension can relate to both stress tolerance *and* competitive strategy, which can blur the relationship between traits and environmental stress gradients, such as that represented by shore height.

Although we examined species’ trait values in relation to their mean shore position, these species often exist in mixed assemblages. Community weighted means allow integration across species in these mixed assemblages (most strongly reflecting the dominants) and indicate consequences for ecosystem functioning (“mass-ratio hypothesis”, Grime 1998). At Bracelet Bay and Oxwich, both at the middle of the exposure gradient, we observed a decline in CWMs of PC1. This means that communities in the more environmentally stressful higher shore were dominated by species with a more conservative thallus structure (low STA/high TDMC). This is consistent with our expectation, with species dominating the higher shore needing to conserve resources (especially water). At Oxwich, there was also a decline in PC2, illustrating that upper shore communities invested proportionally less in holdfasts and were shorter (less competitive). The implications are a slowing of ecosystem functions towards higher on the shore, such as primary productivity (Reich 2014), potentially affecting ecosystem services by reducing the volume of material available to biomass harvesting or fisheries, as well as decreasing the potential for wave attenuation (Smale et al. 2013). Nevertheless, these higher shore communities may still provide valuable functions. For instance, they support and shade faunal assemblages where the threat of desiccation and excessive heat is most extreme. Additionally, their more recalcitrant tissue should persist longer in the environment, driving long-term carbon sequestration (Chung et al. 2011).

However, at sites located at the extremes of the wave exposure gradient, there was no clear relationship between tidal height and CWMs for either PCs. At the very exposed Rhossili, low-shore *L. digitata* appeared to benefit from the level of exposure, as well as the presence of rockpools, and expanded to almost 30% of the shore height, raising CWMs through the mid-shore. At sheltered Angle Bay, the dominance of the outlier *A. nodosum* in many of the quadrats throughout the mid-shore depressed PC1 and elevated PC2. Together, these results show that the correspondence between shore height and CWMs can be disrupted where species show diffuse zonation and shifts in vertical position, or where species possess unusual traits relative to their shore height *– A. nodusum* in this case.

We also aimed to compare within-to between-species variability. Among individuals of the same species, we observed a proportionally small variability across the four sites. Still, this intraspecific variation in traits at such a small spatial scale supports previous studies in LBM (Sjøtun and Fredriksen 1995; Blanchette 1997; Wing et al. 2007) and exemplifies the potential for these species to change their morphology to match environmental conditions, which can be attributed to both adaptive variation and phenotypic plasticity (King et al. 2017). At the interspecific level, we show that considering LBM as a single “functional group”, and therefore ecologically similar, can obscure and underestimate functional diversity. Using multiple continuous functional traits, as demonstrated in this study, provides a solution by allowing individuals and species to be positioned in continuous trait space. Following efforts in terrestrial plants (Díaz et al. 2016), the functional diversity of seaweed should be resolved more generally by expanding this approach to other species of LBM worldwide through coordinated trait screening initiatives. This could shed new light on topics in seaweed ecology, such as their functional biogeography and response to global change drivers (Harley et al. 2012; Violle et al. 2014). Our finding that interspecific trait variability overwhelmed intraspecific variability in LBM supports an initial application of these traits at the species level. Nevertheless, the prevalent, albeit relatively weak, species-site interactions we found here indicate more accurate estimations of functional diversity could ultimately be gained by integrating intraspecific variation.

Our approach has some limitations associated with the scope and focus of our study. We pragmatically used indirect measurements of the gradients of wave exposure (wave fetch of different sites) and tidal emersion (relative shore height), and did not consider direct mediators of stress on individuals such as drag forces or evaporation potential. This is a first step towards understanding spatial variation in seaweed functional traits and is analogous, for example, to using altitude in terrestrial studies of plant functional diversity. We also chose traits as functional markers (*sensu* Garnier et al. 2004), which provided an efficient way to screen multiple dimensions of variability at the cost of fine-scale mechanistic insight. Finally, although not a requisite for functional trait-focused studies (de Bello et al. 2015), future studies may wish to integrate trait and phylogenetic perspectives to formally investigate the evolutionary basis of functional trait diversity among macroalgae, especially when considering more species, and from different clades.

In conclusion, we have reported the first attempt to position species of large brown macroalgae in a continuous, multidimensional, trait space. Reminiscent of findings from terrestrial plants, species were concentrated along a two-dimensional plane defined by axes of resource acquisition and competitive dominance. We found that predicting seaweed trait responses to intertidal (stress) emersion gradient is challenging because 1) resource conservation traits can be related to competitive as well as stress-tolerance strategies, and 2) community zonation patterns can be modified by wave exposure. Overall, our study demonstrates the potential for functional traits to reveal new dimensions of diversity among macroalgae and unify ecological methods and perspectives across ecosystems and evolutionarily divergent producer groups.

## Acknowledgements

This research has been funded by the Brazilian National Council for Scientific and Technological Development (CNPq – Science Without Borders grant 202032/2015-9). We thank Andrew J. King for the contributions to the manuscript; Tom P. Fairchild for the technical help; Mike Burrows for the help obtaining the fetch data; Olga Koppel and Josh Mutter for their help in the field and the lab; Igor S. Pessi for the suggestions to the manuscript and help with the images; and Amanda Cappelatti for the illustrations.

## Author contributions

LC and JG conceived the ideas and designed the methodology; LC collected and analysed the data; LC and JG led the writing of the manuscript; AM contributed throughout the development of the study and the manuscript. All authors contributed critically to the drafts and gave final approval for publication.

## Supplementary material

Figure S1. Boxplots of species traits (log-transformed) across the four study sites. Appendix S1 and Table S1. Description and results of water analysis of the study sites.

Appendix S2, Table S2 and Figure S2. Description and summary of imputation method for missing trait values, and PCA plot after removing samples with missing values.

Appendix S3 and Table S3. Description and summary of residual analysis to test for the effect of tidal height on species’ traits.

Table S4. PERMANOVA including all six traits, with species*site for the species found at more than one site.

Table S5. Mean and standard deviation of relative height where each species was found.

