## Appendix 1 for "Applying continuous functional traits to large brown macroalgae: variation across tidal emersion and wave exposure gradients"

**Supplementary material 1**

**Appendix S1.**

*Water nutrient analysis*: We collected three 50 mL samples of seawater at each site, on three occasions (September 2017, April and July 2018). Samples were filtered and frozen prior to analysis (XY2 Autosampler, Seal Analytical Ltd., Southampton, UK). The results are shown in Table S1.

*Water salinity*: We collected samples for salinity measurement in August 2018. The aim of this analysis was to confirm that the site located in the ria (Angle Bay, Pembrokeshire), into which several rivers flow, had salinity values representative of fully marine conditions and comparable to those of Gower sites. On the same day, six replicates were collected at Angle Bay and six at Oxwich (a site chosen to represent the Gower sites as it is the mid-site geographically) and analysed with a portable probe (Hanna Instruments, Woonsocket, RI, US). Results are given in Table S1.

*Water temperature*: We did not expect any differences in water temperature at these sites due to their similar latitudes. Nevertheless, to confirm this, during the first 10 days of October 2018, we deployed temperature loggers (HOBO Pendant, Onset Computer Co., Bourne, MA, US) at each study site on exposed rock on the middle shore and away from the seaweed canopy. Values were averaged from measurements taken at 10-minute intervals during four hours of daytime high tide. Results are shown in Table S1.

**Table S1**. Averages of seawater nutrient concentrations, salinity, and temperature. TON: Total Oxidized Nitrogen. Standard deviations are given in parenthesis.

| **Sites** | **Nutrients (µmol L^-1^)** | | | | | **Salinity (ppt)** | **Water temp. (^o^C)** |
| --- | --- | --- | --- | --- | --- | --- | --- |
|  | **TON** | **NH_4_** | **NO_2_** | **PO_4_** | **SiO_3_** |  |  |
| Angle Bay | 3.02 (1.9) | 2.98 (1.3) | 0.21 (0.3) | 1.37 (0.4) | 5.77 (3.8) | 33.4 (0.1) | 20.4 (1.3) |
| Bracelet Bay | 3.77 (3.2) | 3.10 (2.3) | 0.24 (0.4) | 1.30 (0.3) | 4.41 (5.7) |  | 20.1 (0.3) |
| Oxwich | 1.44 (0.2) | 3.23 (1.9) | 0.23 (0.3) | 1.18 (0.3) | 5.35 (6.7) | 32.22 (2.4) | 19.9 (0.4) |
| Rhossili | 3.22 (2.4) | 3.98 (2.4) | 0.23 (0.3) | 1.53 (0.4) | 3.90 (3.5) |  | 20.7 (1.3) |
