## Appendix 2 for "Applying continuous functional traits to large brown macroalgae: variation across tidal emersion and wave exposure gradients"

**Supplementary material 2**

**Appendix S2.**

*Imputation of missing trait values.* Of the 167 samples, some had missing values for a trait, which were imputed so we could perform the ordination with all the traits without removing entire samples due to one or two missing values. The total missing values for each trait were: STA (n=4), SAV (35), thickness (2) and holdfast ratio (17). To make sure this imputation did not affect our conclusions, we ran another PCA after removing individuals with missing values instead of imputing them. The contributions of each trait to the first three axes are shown in Table S2, together with contributions of traits with imputed values. The only change was a shift in the order of importance in the loadings of PC2 from holdfast ratio to length. Fig. S2 shows the PCA plot with the samples with missing values removed.

**Table S2.** Contributions of each trait to the first three PCA axes, on all eight study species. On the left, as used for the PCA on main text; on the left, after rows were removed when there was a missing value for one or more traits. Variation explained is given below each axis, in italic. Values are percentages. Main contributions highlighted in bold.

|  | **Missing values imputed** | | | **Missing values removed** | | |
| --- | --- | --- | --- | --- | --- | --- |
|  | Axis 1 | Axis 2 | Axis 3 | Axis 1 | Axis 2 | Axis 3 |
|  | *46.1* | *21.5* | *16.7* | *48.4* | *21.6* | *16* |
| Thickness | 15.99 | **26.41** | 1.03 | 14.05 | **29.87** | 2.62 |
| TDMC | **22.08** | 12.27 | 0.26 | **23.3** | 7.75 | 0.23 |
| STA | **35.47** | 0.04 | 0.00 | **32.43** | 0.01 | 0.04 |
| Length | 4.73 | 17.97 | **50.75** | 4.3 | **27.2** | **45.9** |
| SAV | **17.79** | 15.74 | 4.06 | **19.98** | 12.85 | 2.15 |
| Holdfast ratio | 3.94 | 27.55 | **43.88** | 5.9 | 22.3 | **49.05** |

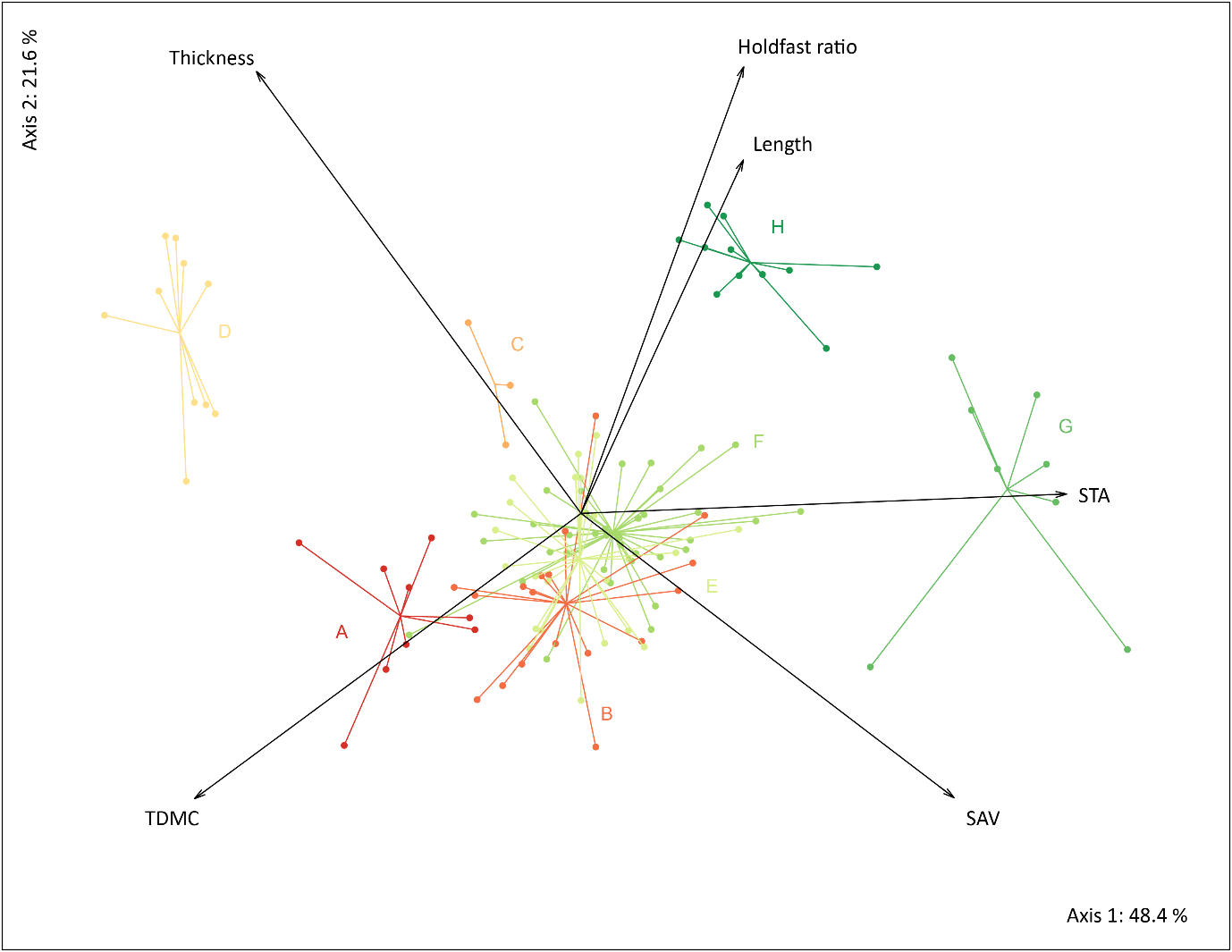

**Figure S2**. First two PCA axes (samples with missing values removed) of large brown macroalgae species and functional traits at all study sites in south Wales, UK. All eight sampled species are included, and, with the exception of *H. siliquosa* and *S. latissima* were found in at least three of the four sites (see main text in *Methods* and *Results* for more information on species occurrences and use in analyses). A: *Pelvetia canaliculata*; B: *Fucus spiralis*; C: *Halidrys siliquosa*; D: *Ascophyllum nodosum*; E: *F. vesiculosus*; F: *F. serratus*; G: *Saccharina latissima*; and H: *Laminaria digitata*.
