## Appendix 3 for "Applying continuous functional traits to large brown macroalgae: variation across tidal emersion and wave exposure gradients"

**Supplementary material 3**

**Appendix S3.**

*Species traits as response to tidal height.* To further investigate intraspecific variability, for each trait we checked if, after accounting for site and species differences (ANOVA with site and species interaction), residual variability could be attributed to variation in tidal height of collected samples. The results are shown in Table S3.

**Table S3.** Summary of analysis with residuals of each trait from site*species analysis, with relative height as independent variable.

| **Species** | **Estimate** | **Std. error** | **T value** | **P value** |
| --- | --- | --- | --- | --- |
| Thickness | 0.00 | 0.05 | 0.15 | 0.87 |
| TDMC | 0.06 | 0.06 | 0.99 | 0.32 |
| STA | -0.04 | 0.07 | -0.5 | 0.6 |
| Length | 0.02 | 0.17 | 0.09 | 0.92 |
| SAV | 0.06 | 0.11 | 0.55 | 0.58 |
| Holdfast ratio | -0.29 | 0.29 | -1.02 | 0.31 |
