## Supplemental figure 1 for "Applying continuous functional traits to large brown macroalgae: variation across tidal emersion and wave exposure gradients"

**Supplementary material**

**Figure S1**

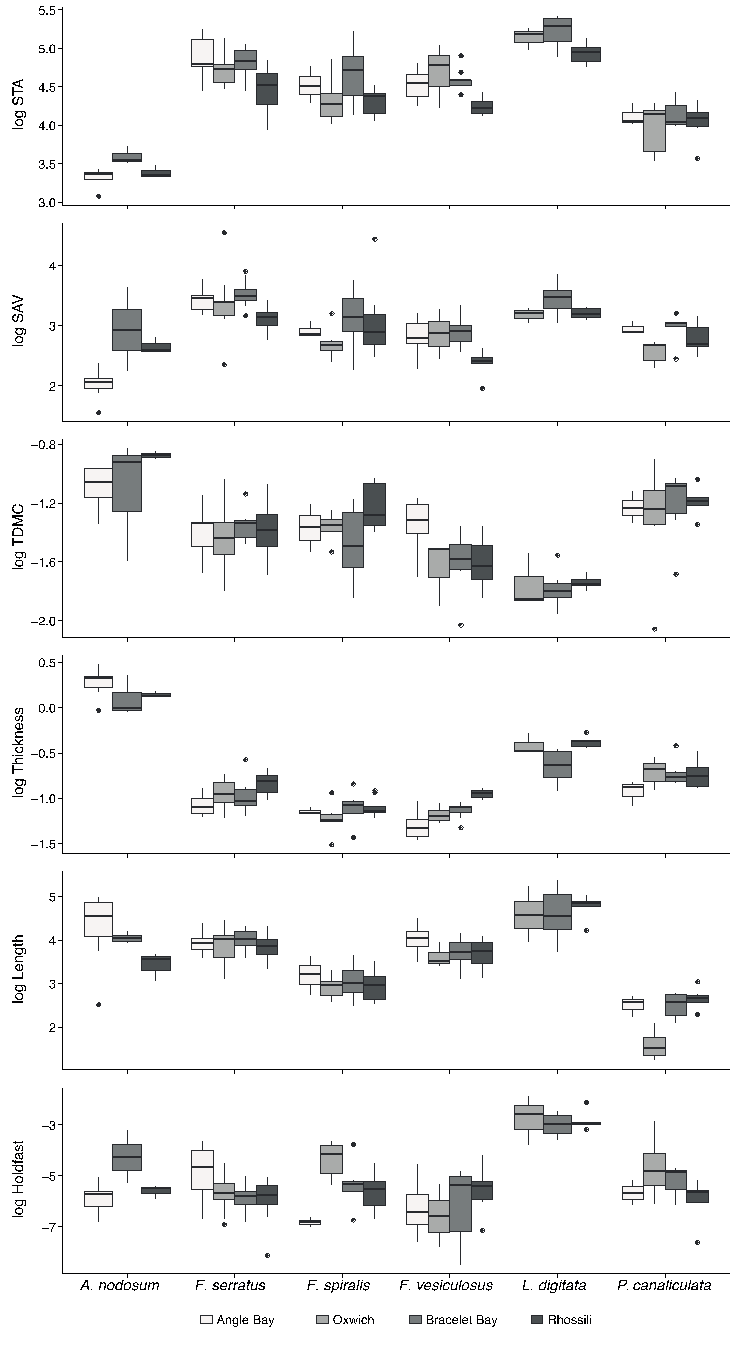

**Figure S1.** Boxplots of species traits (log-transformed) across the four study sites, for the six species found at more than one site. Grey hue indicates intensity of wave exposure.
