## Supplemental table 4 for "Applying continuous functional traits to large brown macroalgae: variation across tidal emersion and wave exposure gradients"

**Supplementary material 4**

Table S4. Result of the PERMANOVA including all six traits, individual level, for the six species that occurred at more than one site. Degrees of freedom (df), and test statistics (sum of squared differences, mean of squared differences, F-value and P value) are given, as well as an estimated variance explained (Partial R^2^).

|  | **df** | **Sum sq** | **Mean SS** | **F** | **R^2^** | **P value** |
| --- | --- | --- | --- | --- | --- | --- |
| Site | 3 | 31.92 | 10.64 | 9.61 | 0.067 | **0.001** |
| Species | 5 | 251.09 | 50.22 | 45.69 | 0.529 | **0.001** |
| Site * Species | 13 | 44.57 | 3.43 | 3.12 | 0.094 | **0.001** |
| Residuals | 134 | 147.29 | 1.10 |  | 0.310 |  |
