## Supplemental table 5 for "Applying continuous functional traits to large brown macroalgae: variation across tidal emersion and wave exposure gradients"

**Supplementary material 5**

**Table S5.** Mean and standard deviation of relative height (RH) where each species occurred, at all four study sites, from the 2017 trait survey and the 2018 abundance survey.

| **Species** | **Mean (SD) RH** | |
| --- | --- | --- |
|  | **Trait survey** | **Abundance survey** |
| *Ascophyllum nodosum* | 0.49 (0.12) | 0.45 (0.12) |
| *Fucus serratus* | 0.35 (0.15) | 0.26 (0.12) |
| *Fucus spiralis* | 0.71 (0.05) | 0.64 (0.11) |
| *Fucus vesiculosus* | 0.45 (0.15) | 0.43 (0.10) |
| *Halidrys siliquosa* | 0.58 (0.00) | NA |
| *Laminaria digitata* | 0.11 (0.04) | 0.21 (0.10) |
| *Pelvetia canaliculata* | 0.75 (0.05) | 0.74 (0.04) |
| *Saccharina latissima* | 0.30 (0.00) | 0.31 (0.00) |
